# Regulation of bacterial Type III Secretion System export gate opening

**DOI:** 10.1101/2021.01.17.426956

**Authors:** Owain J. Bryant, Gillian M. Fraser

## Abstract

Type III Secretion Systems (T3SS) transport proteins from the bacterial cytosol for assembly into cell surface nanomachines or for direct delivery into target eukaryotic cells. At the core of the flagellar T3SS, the FlhAB-FliPQR export gate regulates protein entry into the export channel whilst maintaining the integrity of the cell membrane. Here, we identify critical residues in the export gate FliR plug that stabilise the closed conformation, preserving the membrane permeability barrier, and we show that the gate opens and closes in response to export substrate availability. Our data indicate that FlhAB-FliPQR gate opening, which is triggered by substrate export signals, is energised by FlhA in a proton motive force-dependent manner. We present evidence that the export substrate and the FliJ stalk of the flagellar ATPase provide mechanistically distinct, non-redundant gate-activating signals that are critical for efficient export.

## Introduction

Type III Secretion Systems (T3SS) transport proteins across the bacterial inner and outer membranes and, in the case of the virulence T3SS (vT3SS), across the plasma membrane of target eukaryotic cells ^1,2,3^. The flagellar T3SS (fT3SS) is required for assembly of rotary flagella, which facilitate cell motility^1,2,3,4^. Despite their different functions, the vT3SS and fT3SS are evolutionarily related and contain conserved core export components^5,6,7,8,9,10,11^.

For construction of the bacterial flagellum, most of the flagellar components are exported by the fT3SS machinery, which is embedded in the cell envelope. The core of the export machinery is made up of five highly conserved proteins (FlhA, FlhB, FliP, FliQ and FliR). FlhA forms a nonameric ring structure comprising a cytoplasmic domain which functions as a docking platform for export cargo and an N-terminal region (FlhA_N_) with putative proton conducting activity^5,12 13,14^. Recent cryo-ET data has shown that FlhA_N_ wraps around the base of a helical-dome composed of FliPQR and the N-terminal region of FlhB (FlhB_N_)^13^. Together, FlhA and PQRB_N_ constitute the ABPQR export gate, which regulates entry of flagellar subunits into the central channel that runs the length of the nascent flagellum^5,12,13^. In addition to the ABPQR export gate, an ATPase complex (FliHIJ) located in the cytoplasm converts the fT3SS into a highly efficient pmf-driven protein export machine^10,15,16,17^.

Recent structural studies of FliPQR and FlhB_N_ have shown that multiple interactions stabilise the closed export gate ^5,6,12,18^. A central plug formed by residues 106-120 of FliR, a methionine rich loop (M-gate) within FliP, and ionic interactions between adjacent FliQ subunits maintain the closed conformation^5,6,18^. Structural analysis of the export gate in the open conformation revealed multiple rearrangements within the M-gate and an upward displacement of the FliR plug, indicating that energy must be provided to open the gate and permit subunit translocation across the inner membrane^18^. We reasoned that, as the PQRB components of the gate are positioned adjacent to the proposed proton conducting FlhA membrane protein, FlhA might use the energy derived from the pmf to promote gate opening. Moreover, we hypothesised that gate opening must be regulated by one or more factors to permit subunit export whilst maintaining the membrane permeability barrier. We set out to test these hypotheses experimentally.

## Results

### Replacement of critical residues in the FliR plug destabilises the closed conformation of the fT3SS export gate

The fT3SS ABPQR export gate resides in a closed conformation in the absence of subunit export^5,6,13,11,18^. We set out to engineer the export gate to destabilise the closed conformation and favour the open state, reasoning that mutation of contact points between adjacent gate subunits would reduce the energy barrier required for gate opening (Fig.1A). To screen for potential gate opening mutants, we used a subunit variant (FlgD_short_) that we have previously shown is unable to trigger efficient opening of the export gate^19^. We predicted that mutations which promoted export gate opening would also supress the FlgD_short_ motility defect. Recent structural information has revealed the potential gating mechanism of the FliPQR gate components. A region of FliR (residues 106-121) forms a plug that occludes the gate channel, and this plug must be displaced to achieve gate opening^18^. In the FliR plug, residues Phe-110 and Phe-113 make multiple contacts within the core of the export gate, and might have a central role in gate opening^5,12^. To assess the importance of selected FliR residues in gate opening, we replaced Phe-110 and/or Phe-113 with alanine, and replaced Gly-117 with a bulkier charged aspartate residue and screened for suppression of the FlgD_short_ motility defect (Fig. 1). We found that all three of these FliR-plug mutations (F_110_A, F_113_A and G_117_D) supressed the FlgD_short_ motility defect, yet did not increase export or motility of cells encoding wild type FlgD (Fig. 1B, 1C and S1, S2), indicating these mutations destabilise the closed export gate conformation. The FliR-G_117_D mutation was previously proposed to rescue an interaction between FliR and FlhA, but the data presented here indicate that the G_117_D, F_113_A and F_115_A mutations destabilise the closed conformation of the export gate (Fig. 1D)^20^.

**Figure 1.**
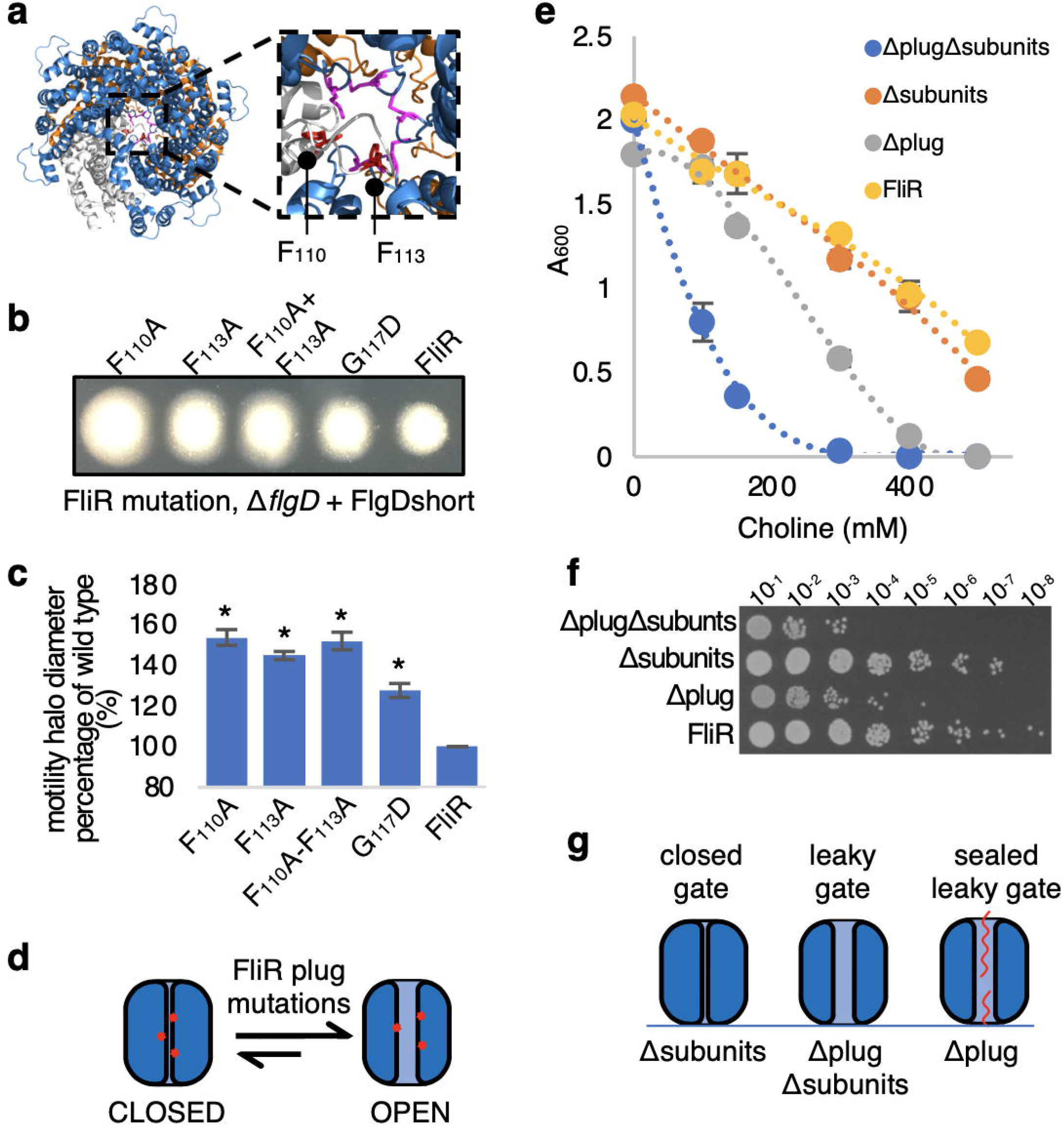
Mutations within the FliR plug destabilise the closed export gate conformation. **a**. Top view of the cryo-EM structure of the closed conformation of the FliPQR export gate complex (PDB:6F2D). Phenylalanine residues 110 and 113 of FliR (grey) are displayed in red. Methionine residue 210 of FliP (blue) is displayed in magneta. FliQ is displayed in orange. **b**. Swimming motility of recombinant *Salmonella flgD* null strains producing chromosomally-encoded FliR variants (F_110_A, F_113_A, F_110_A+F_113_A or G_117_D) or wild type FliR (FliR). Strains produced a pTrc99a plasmid-encoded FlgD subunit variant in which residues 9-32 were replaced with two repeats of the six amino acid sequence Gly-Ser-Thr-Asn-Ala-Ser (FlgD_short_). Motility was assessed in 0.25% soft-tryptone agar containing 100 μg/ml ampicillin and 50 μM IPTG and incubated at 37°C for 16 hours. **c**. The mean motility halo diameter of recombinant *Salmonella flgD* null strains producing chromosomally-encoded FliR variants (F_110_A, F_113_A, F_110_A+F_113_A or G_117_D) or wild type FliR (FliR) and expressing the FlgD_short_ variant was plotted as a percentage of the wild type FliR strain (FliR) producing FlgD_short_ (right hand bar). Error bars represent the standard error of the mean calculated from at three biological replicates. * indicates a *p*-value < 0.05. **d**. A cartoon schematic illustrating the effect of FliR plug mutations (red) on the conformation of the export gate complex (blue). Data indicates that FliR plug mutations destabilise the closed export gate conformation (left) and bias the open conformation (right). **e**. Choline sensitivity of a *Salmonella* strain producing a chromosomally-encoded FliR variant in which residues 110-116 of the FliR plug is deleted (Δplug, grey), a *Salmonella* strain deleted for the genes (*flgBCDEFGHIJ*) that encode most of the early flagellar rod and hook subunits (Δsubunits, orange), a *Salmonella* strain that both produces the chromosomally-encoded FliRΔplug variant and is deleted for the genes (*flgBCDEFGHIJ*) that encode most of the early flagellar rod and hook subunits (ΔplugΔsubunits, blue) or a wild type *Salmonella* strain (FliR, yellow). Cells were grown in Terrific broth containing varying concentrations of choline (as indicated) and the optical density (A_600_) measured following 4.5 hours. **f**. Serial dilutions of the above *Salmonella* strains following 4.5 hours of growth in Terrific broth supplemented with 300 mM choline were spotted onto LB agar plates using the Miles and Misra method. The strains are indicated on the left and the density of the inoculum is indicated along the top. **g**. A cartoon schematic illustrating the effect of the FliR plug deletion on the gating activity of the export gate complex (blue). Strains that produce fewer subunits (Δsubunits and ΔplugΔsubunits) have fewer available subunits (red) for export to seal the leaky phenotype of the export gate containing the FliR plug deletion (Δplug, right).

We hypothesised that deletion of the FliR plug would promote the open gate conformation and, as a result, increase the permeability of the inner membrane. Bacterial cells that fail to maintain the permeability barrier across the inner membrane are more sensitive to a number of chemical agents, such as choline^21^. To determine whether a strain encoding a FliR plug deletion variant (FliRΔ110-116, hereafter termed FliRΔplug) was sensitive to choline, we performed growth inhibition assays by culturing cells in media containing increasing concentrations of choline (Fig 1E). We found that the FliRΔplug strain was sensitised to choline compared to wild type *Salmonella*, indicating that the FliR plug is important for formation of a tight barrier that is impermeable to small molecules, and for maintaining the closed conformation of the fT3SS export gate (Fig. 1E, 1F, S3).

### The export gate fluctuates between open and closed states in response to substrate availability

A previous study showed the M_210_A mutation in the FliP component of the export gate sensitised cells to a variety of chemical agents^21^. This sensitivity could be reversed by trapping a subunit in the export channel, suggesting that the export gate can form an effective seal around a substrate during transit^21^. We reasoned that if the export gate fluctuates between open and closed conformations in response to the availability of substrate, strains producing fewer flagellar subunits would, at any given time, have fewer subunits transiting through the fT3SS to seal the leaky FliRΔplug gate. To test this, we constructed a strain deleted for the genes that encode most of the early flagellar subunits by replacing a large portion of the *flg* operon (*flgB, flgC, flgD, flgE, flgF, flgG, flgH, flgI* and *flgJ*) with a kanamycin resistance cassette. In addition, we constructed a second strain in which the FliR plug and the rod/hook genes were deleted. This FliRΔplug-Δsubunit strain was significantly more sensitive to choline than the FliRΔplug strain, indicating that actively transiting subunits partially seal the export gate when the FliR plug is absent (Fig. 1E, 1F, S3). Notably, the Δsubunit strain (which produces wild type FliR) was not more sensitive to choline than wild type *Salmonella*, indicating that decreased subunit availability does not increase the permeability of the wild type export gate to small molecules (Fig. 1E, 1F, 1G). This suggests that the export gate must fluctuate between open and tightly closed conformations, in response to availability of subunits at the export machinery.

### Destabilising the closed conformation of the export gate alleviates the export defect associated with a variant of the gate component FlhA

Opening of the export gate is thought to be energised, in part, by harnessing of the pmf by FlhA. A previous study proposed that FlhA interacts with FliR, as mutations in *fliR* were found to suppress the motility defect of a weakly motile *Salmonella* (Δ*fliH-fliI, flhB*-P_28_T, *flhA*-K_203_W) that contained deletions in flagellar ATPase genes (*fliI* and *fliH*), a suppressor mutation in *flhB*, P_28_T, which overcomes the loss of FliH and FliI, and a mutation in *flhA*, K_203_W, that attenuates motility and export ^20^. However, structural studies subsequently revealed that these FliR suppressor mutations are located in the core of the FliPQR complex, indicating that this region of FliR is unable to contact FlhA within the assembled fT3SS^5^. We have shown that one of these suppressor mutations in the FliR plug (FliR-G_117_D) destabilises the closed conformation of the export gate (Fig. 1B and 1C). Based on these data, we hypothesised that FliR-G_117_D might suppress the motility and export defects associated with mutation of FlhA-K_203_ by destabilising the closed conformation of the export gate, rather than by restoring an interaction between FliR and FlhA. To test this, we assessed whether the export and motility of strains encoding FlhA-K_203_A could be recovered by introducing mutations (FliR-F_113_A or FliR-G_117_D) that destabilise the closed gate conformation. We found that, as predicted, FliR-F_113_A and FliR-G_117_D suppressed the export and motility defects associated with FlhA K_203_A (Fig. 2). The data indicate that the *flhA*-K_203_A strain is unable to open the export gate efficiently and that introduction of *fliR* mutations that destabilise the closed gate might lower the energy barrier that must be overcome by FlhA-K_203_A to open the gate (Fig. 2A-2B).

**Figure 2.**
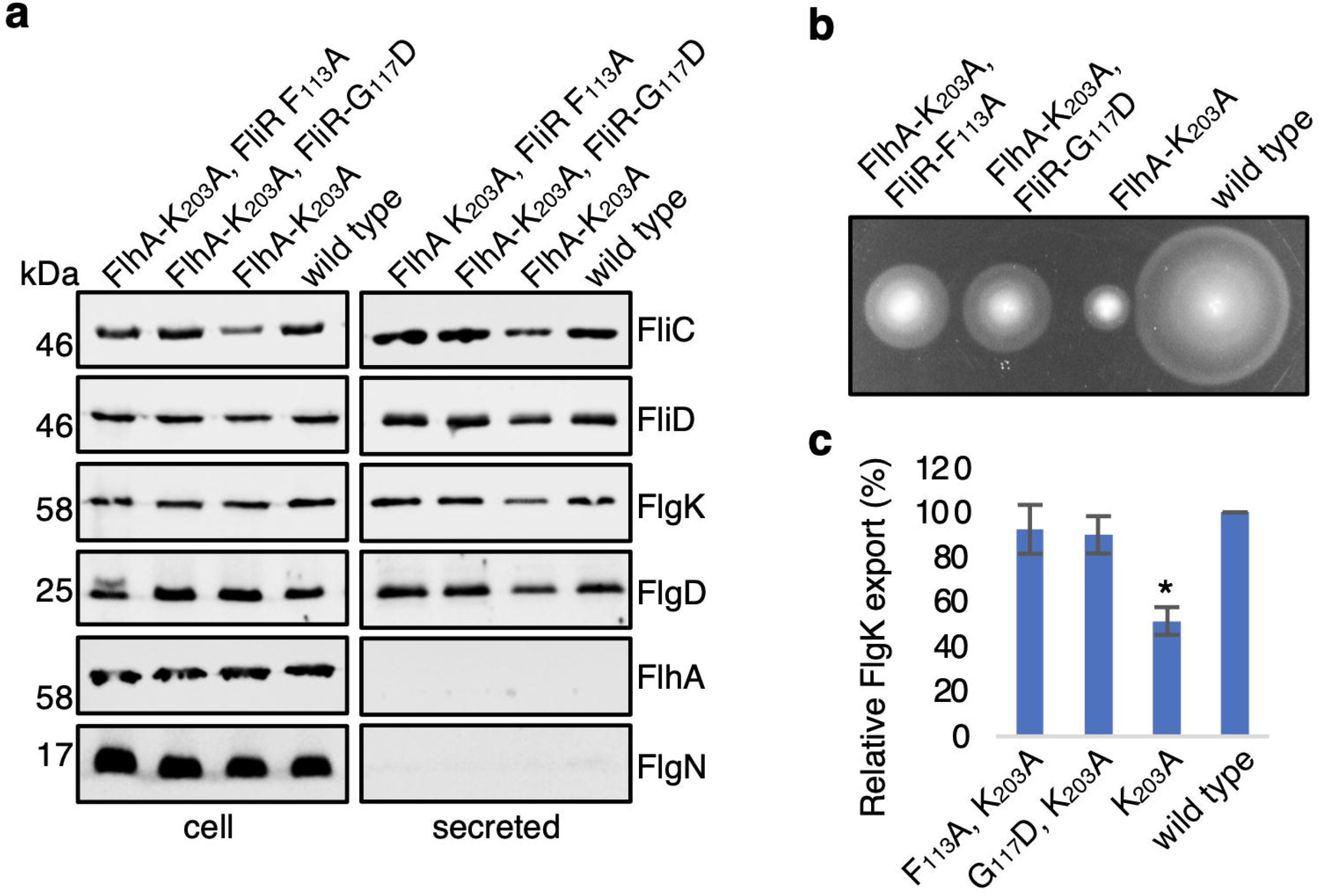
Mutations in the FliR plug suppress the FlhA-K_203_A motility and export defects. **a**. Whole cell (cell) and supernatant (secreted) proteins from late exponential-phase cultures of a wild type Salmonella strain (wild type), a *Salmonella* strain producing a chromosomally-encoded FlhA-K_203_A variant (FlhA K_203_A) or *Salmonella* strains producing both a chromosomally-encoded FlhA-K_203_A variant and chromosomally encoded FliR variants (F_113_A or G_117_D). Proteins were separated by SDS (15%)-PAGE and analysed by immunoblotting with anti-FliC, anti-FliD, anti-FlgK, anti-FlgD, anti-FlhA or anti-FlgN polyclonal antisera. Apparent molecular weights are in kilodaltons (kDa). **b**. Swimming motility (0.25% soft tryptone agar) of the same strains were carried out at 37°C for 4-6 hours. **c**. Levels of FlgK in culture supernatants from Immunoblots were quantified using ImageStudiolite and plotted as a percentage of FlgK exported by the wild type strain. Error bars represent the standard error of the mean calculated from three biological replicates. * indicates a *p*-value < 0.05.

### Increasing the pmf suppresses the FlhA-K_203_A export defect

Our results support the view that FlhA has a role in opening the export gate and that FlhA-K_203_ is critical for efficient gate opening. As FlhA has been proposed to facilitate subunit export by functioning as a proton-conducting channel^14,17,20^, we hypothesised that the FlhA-K_203_A mutation might render the export machinery unable to couple pmf to efficient gate opening. If this were the case, the gate opening and subunit export defects of a strain carrying chromosomally encoded *flhA-K*_*203*_*A* might be overcome by increasing the pmf. To test this, we artificially increased the pmf across the bacterial inner membrane by driving intracellular proton consumption^22^. Cells supplemented with arginine convert intracellular protons and arginine to agmatine and CO_2_ through the action of the cytoplasmic enzyme arginine decarboxylase, which effectively acts as a proton sink in the cytosol and therefore increases the pmf^22^ (Fig. 3A). We performed a modified subunit export assay in which cells were grown to mid-exponential phase, washed, resuspended in media containing 20 mM arginine and culture supernatants were collected after 30 mins. Addition of arginine to cultures of the *flhA*-K_203_A strain increased subunit export by 40%, whereas subunit export in the wild type strain was unchanged (Fig. 3B). Deletion of the two arginine decarboxylase enzymes (*speA* and *adiA*) encoded by *Salmonella* abolished the arginine-dependent increase in subunit export by the *flhA*-K_203_A strain, indicating that the FlhA-K_203_A mutation prevented efficient use of the pmf by the export machinery. The data are consistent with wild type FlhA utilising the pmf to energise export gate opening (Fig. S4).

**Figure 3.**
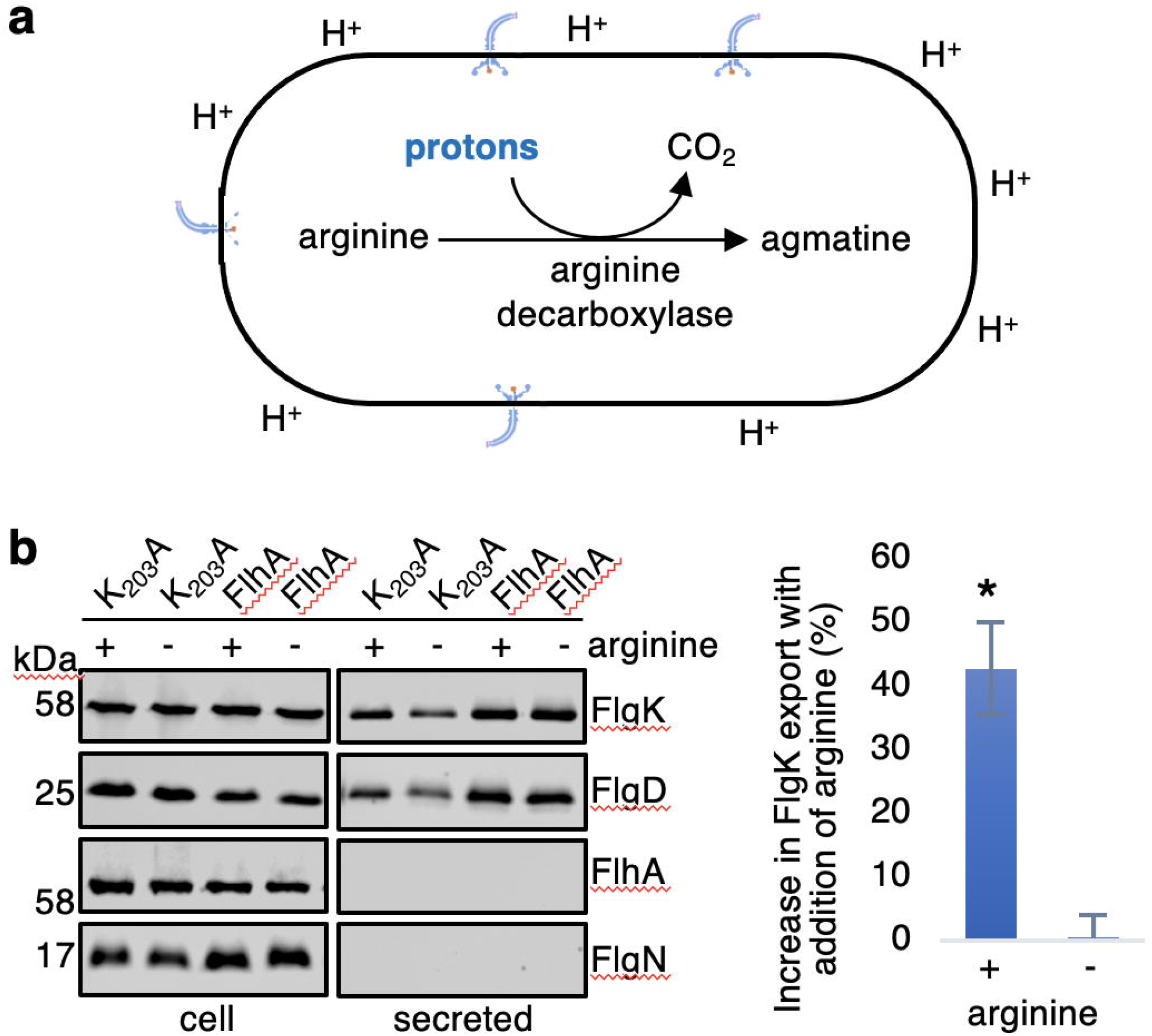
Increasing the proton-motive force suppresses the *flhA*-K_203_A export defect. **a**. A cartoon schematic of a cell containing arginine decarboxylase enzyme which catalyses the conversion of L-arginine into agmatine and carbon dioxide (CO_2_). The arginine decarboxylation reaction consumes a proton (H^+^) from the cell cytoplasm, reducing the number of protons in the cell, effectively increasing the proton-motive force. **b**. Whole cell (cell) and supernatant (secreted) proteins from late exponential-phase cultures of a wild type *Salmonella* strain (FlhA) or a *Salmonella* strain producing a chromosomally-encoded FlhA-K_203_A variant (K_203_A). Cells were supplemented with (+) or not supplemented with (-) 20 mM arginine 30 minutes prior to collection of whole cell and culture supernatants. Proteins were separated by SDS (15%)-PAGE and analysed by immunoblotting with anti-FlgK, anti-FlgD, anti-FlhA or anti-FlgN polyclonal antisera (Left). Apparent molecular weights are in kilodaltons (kDa). Levels of FlgK in culture supernatants from Immunoblots for the FlhA-K_203_A strain were quantified using ImageStudioLite and plotted as a percentage of FlgK exported by strains supplemented with arginine (+) to strains not supplemented with arginine (-) (right). Error bars represent the standard error of the mean calculated from three biological replicates. * indicates a *p*-value < 0.05.

### Two distinct non-redundant signals activate the pmf-driven export machinery

We have found that the N-terminus of rod/hook subunits contains a hydrophobic signal that triggers opening of the export gate^19^, a process that would require an input of energy from either the flagellar ATPase and/or the proton motive force. We reasoned that the subunit ‘gate opening’ signal must somehow be transmitted to FlhA, triggering it to energise opening of the export gate. Previous studies have shown that, in addition, interaction of FlhA with the FliJ stalk component of the ATPase is required to convert the fT3SS into a highly efficient ΔΨ driven export machine, suggesting that the FliJ-FlhA interaction might be a second signal required for export gate opening^17^. Loss of either the FliJ-FlhA interaction or the subunit N-terminal hydrophobic signal can be overcome by suppressor mutations in the export machinery: FliH-FliI-FliJ loss can be overcome by mutations in FlhA or FlhB^23,24^, and we have shown that loss of the interaction between the FlgD_short_ subunit N-terminal hydrophobic signal and the export machinery can be overcome by mutations in the genes encoding FliP or FliR that destabilise the export gate closed conformation^19^. We now wanted to test whether the suppressor mutations in the ABPQR export gate (FliR-F_113_A or FliR-G_117_D) that recovered export of FlgD_short_ could also recover motility and export in strains deleted for the ATPase genes (in which the FliJ-FlhA is lost; Fig. 4A and 4B). To test this, we constructed strains in which the chromosomal *fliR* gene was replaced with *fliR* alleles containing the gate opening mutations (F_113_A or G_117_D) in combination with deletions in the genes that encode the ATPase components FliH and FliI. These strains were also deleted for the gene encoding the flagellar anti-sigma factor FlgM, which is exported during the filament stage of assembly^25^. Deletion of *flgM* ensures that flagellar gene expression is not influenced by any differences in export efficiency between strains^26^ (Fig. 4). Mutations that promoted the open gate conformation (FliR-F_113_A or FliR-G_117_D) did not recover motility and export in strains deleted for the ATPase complex (Fig. 4A and 4B). This indicates that mutations that promote the open conformation of the FliPQR gate cannot bypass the loss of FliJ-dependent activation of FlhA.

**Figure 4.**
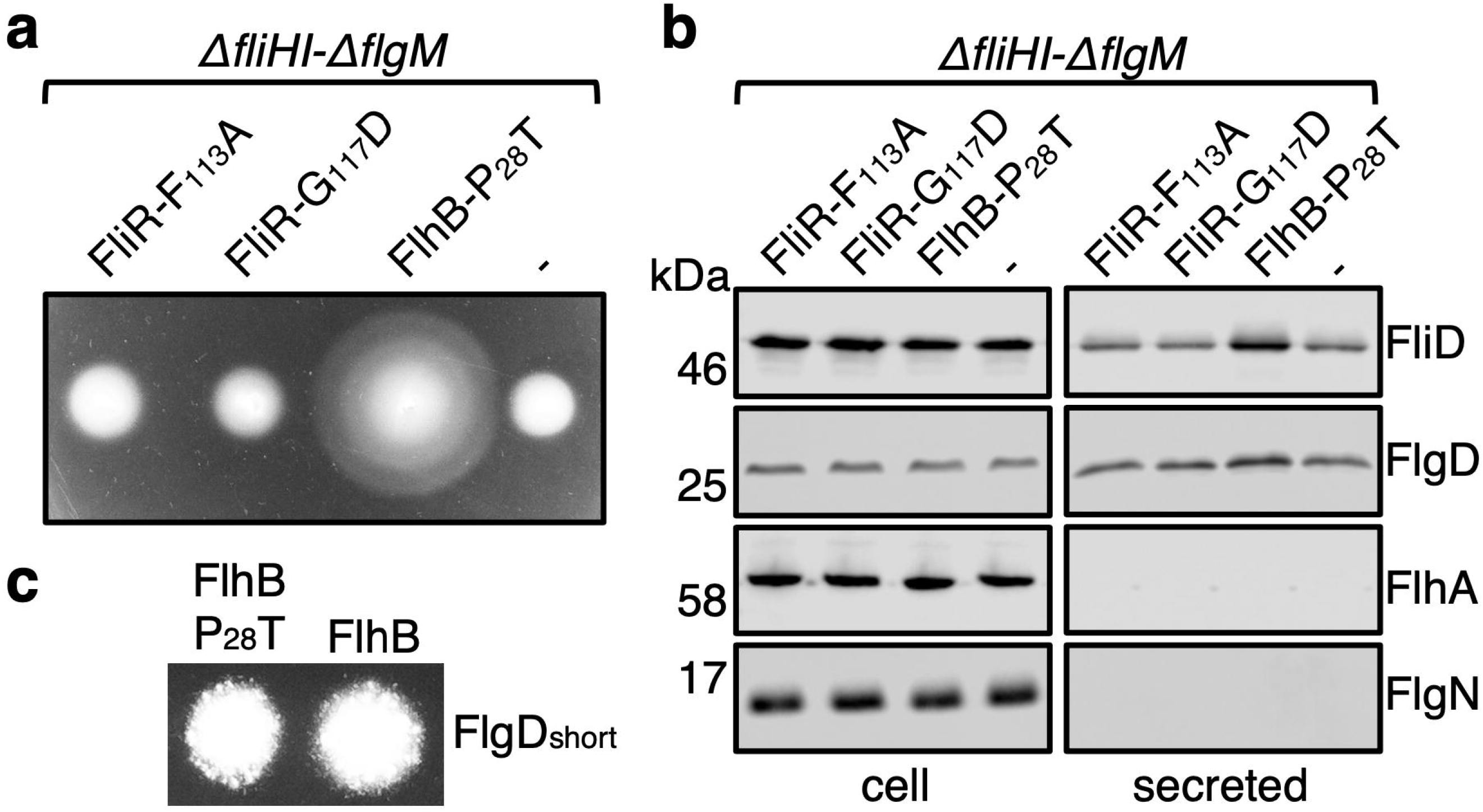
Mutations in the FliR plug do not suppress export and motility defects of strains deleted for the ATPase complex. **a**. Swimming motility of recombinant *Salmonella* strains deleted for the genes that encode; the anti-sigma(28) factor (*flgM*), the flagellar ATPase negative regulator (*fliH*) and the flagellar ATPase subunit (*fliI*) and producing either a chromosomally-encoded FlhB-P_28_T variant or producing chromosomally encoded wild type FliR (-) or its variants (F_113_A or G_117_D). Motility was assessed in 0.25% soft-tryptone agar and incubated at 37°C for 4-6 hours. **b**. Whole cell (cell) and supernatant (secreted) proteins from late exponential-phase cultures of the same strains were separated by SDS (15%)-PAGE and analysed by immunoblotting with anti-FliD, anti-FlgD, anti-FlhA or anti-FlgN polyclonal antisera. Apparent molecular weights are in kilodaltons (kDa). **c**. Swimming motility of recombinant *Salmonella flgD* null strains producing chromosomally-encoded wild type FlhB or its variants (FlhB-P_28_T). Both strains produced a pTrc99a plasmid-encoded FlgD subunit variant in which residues 9-32 were replaced with two repeats of the six amino acid sequence Gly-Ser-Thr-Asn-Ala-Ser (FlgD_short_). Motility was assessed in 0.25% soft-tryptone agar containing 100 μg/ml ampicillin and 50 μM IPTG and incubated at 37°C for 16 hours.

We then assessed whether the FlhB-P_28_T suppressor, which enables bypass of flagellar ATPase activity (and, as a result, bypass of the FliJ-FlhA interaction), could recover efficient gate opening by FlgD_short_ (Fig. 4C). A *Salmonella flgD* null strain encoding *flhB*-P_28_T and producing FlgD_short_ was found to be no more motile than the parental *Salmonella flgD* null strain producing FlgD_short_ (Fig 4C and S5). This indicates that the FlhB-P_28_T mutation does not promote opening of the export gate, but instead restores fT3SS activity in the absence of the FliJ activation signal by an alternative mechanism (Fig. 4C).

These data indicate that mutations in *fliP* or *fliR* that promote the open conformation of the export gate and mutations that bypass the need for the FliJ-FlhA interaction are unable to compensate for each other, indicating that the subunit N-terminal hydrophobic export signal and the FliJ-FlhA interaction signal are non-redundant, *i*.*e*. both signals are required to activate pmf-driven opening of the ABPQR export gate by FlhA (Fig. 5).

## Discussion

Transport of substances across the bacterial inner membrane is highly selective, thereby maintaining the essential electrochemical gradients that drive numerous processes at the cell membrane, including ATP synthesis and protein export. The fT3SS translocates proteins efficiently across the inner membrane without compromising the permeability barrier^5,18,21^. The ABPQR flagellar export gate contains three constriction points that are proposed to maintain the gate in a closed conformation, preserving the integrity of the membrane permeability barrier^5^. Here, we identified point mutant and deletion variants of proteins in the ABPQR export gate that destabilise its closed conformation. We show that either by destabilising the export gate closed conformation or by increasing the pmf, the flagellar export defects of strains encoding FlhA-K_203_A can be alleviated, indicating that FlhA harnesses the pmf to energise opening of the export gate complex. Additional *in vivo* experiments revealed that FlhA requires activation by both the flagellar ATPase stalk component FliJ and an N-terminal hydrophobic signal in the export substrate to trigger efficient gate opening and subunit export.

### Mutational analysis of the FliR-plug identifies variants that destabilise the closed conformation of the export gate

Structures of the FliPQR export gate have revealed that, in the absence of the export substrate, multiple non-covalent interactions in the gate complex stabilise the closed conformation^5,12,18^. In particular, a 15-residue region of FliR forms a plug that occludes the central channel of the gate. Two residues within the FliR-plug, FliR-F_113_ and FliR-F_115_, interact with a concentric ring of conserved methionine residues (M-gate) in FliP^5^. Our mutational analyses identified FliR variants, FliR-F_113_A and FliR-F_115_A, that destabilise the closed conformation of the export gate^5^, indicating that these residues help to maintain the gate in its closed state in the absence of export substrate. An additional FliR-plug variant, FliR-G_117_D, also destabilised the closed gate, suggesting that the introduction of a bulky, charged side chain at this site prevents tight closure of the gate. A recent structure of the open export gate revealed multiple rearrangements within the M-gate and an upward displacement of the FliR plug to allow subunit passage^18^. This indicates that the FliR plug, which occludes the export channel in the closed gate conformation, must be displaced to allow passage of unfolded subunits through the gate. A strain containing a deletion within the FliR-plug (residues 110-116) displayed choline sensitivity, consistent with the plug forming a tight seal to maintain the membrane permeability barrier (Fig 1). Notably, we found that a FliR-plug deletion strain that produced fewer subunits was significantly more sensitive to choline than a strain that encoded wild type FliR and produced fewer subunits (Fig 1). These data indicate that (i) subunit transit through the export gate can partially rescue the choline-sensitive phenotype associated with loss of the FliR-plug, and (ii) that when no subunits available for transit, the wild type export gate closes to maintain the membrane permeability barrier. These observations have important implications for our understanding of how the export gate opens. Sequential rounds of gate opening and closing must occur in response to the presence or absence of export substrates at the membrane export machinery. Similar gating mechanisms have been observed in other protein conducting channels, such as the SecY channel, which similarly contains a constriction formed by ring of hydrophobic residues and a central plug that forms a seal^27^. Deletion of either the central plug or the ring of hydrophobic residues in SecY results in increased permeability of the channel, and electrophysiology experiments showed that these deletions cause the channel to alternate between an open and closed state^28^. Furthermore, molecular dynamic simulations indicate that the force required for opening the gate is reduced in SecY plug deletion mutants^29^. We found that deletion of the FliR plug similarly destabilised the closed conformation of the export gate and concomitantly increased permeability of the inner membrane (Fig. 1).

### FlhA activity is required for efficient opening of the export gate

A previous study identified suppressor mutations in the gene encoding FliR that rescued the export defect of a strain that contained mutations in *flhA* (K_203_W) and *flhB* (P_28_T), as well as deletions in the genes encoding flagellar export ATPase components (*fliH* and *fliI*)^20^. The authors proposed that the FliR suppressor mutations might recover an interaction between FliR and FlhA^20^. One of these suppressor mutations (FliR-G_117_D) is positioned within the core of the FliPQR complex and introduces a bulky, charged side chain into the lumen of the export gate. We showed that the FliR-G_117_D mutation partially suppressed the motility defect associated with a subunit (FlgD_short_) that is unable to trigger efficient opening of the export gate, indicating that the FliR-G_117_D mutation might instead destabilise the export gate’s closed conformation (Fig. 1B and 1C). We hypothesised that by the same mechanism, the FliR-G_117_D mutation might also suppress the motility and export defects associated with FlhA-K_203_. *Salmonella* strains carrying the *flhA*-K_203_A mutation in combination with mutations in *fliR* (F_113_A or G_117_D), which destabilise the closed conformation of the export gate, displayed enhanced flagellar export and motility compared to the parental *Salmonella flhA*-K_203_A strain (Fig. 2). This indicates that FlhA-K_203_A is unable to facilitate efficient export gate opening and, as a result, attenuates subunit export and cell motility.

Cryo-electron tomograms of the *Salmonella* injectisome have revealed that the FlhA homologue, InvA, is positioned adjacent to the homologues of the FliPQR export gate components, SpaPQR^13^. As FlhA is proposed to function as a proton conducting channel that energises export, we reasoned proton translocation might be coupled to gate opening, possibly through transmission of conformational changes in FlhA to the adjacent FliPQR-FlhB components of the export gate^13,14,20^. In support of this view, we found that by increasing the pmf across the inner membrane, through supplementation of cell cultures with arginine, subunit export by the *flhA*-K_203_A strain could be enhanced by 40%, suggesting that inefficient export in the *flhA*-K_203_A strain is caused by its decreased ability to couple the pmf to gate opening (Fig. 3A).

### Two distinct signals activate the pmf-driven export machinery

Previous studies have shown that the FliJ stalk component of the flagellar ATPase interacts with FlhA to convert the export machinery into a highly efficient ΔΨ-driven export machine^17,30,31,32^. The FliH and FliI components of the ATPase facilitate the FliJ-FlhA interaction to promote full activation of the export gate^17,31^. These findings indicate that FliJ binding to FlhA modulates its proton conducting ability, analogous to the interactions between the stalk component of the F_1_ ATPase and the membrane-localised F_0_ channel that drive proton translocation across the bacterial inner membrane. The need for activation of FlhA by the FliJ stalk, to facilitate efficient use of the ΔΨ component of the pmf and energise gate opening, can be bypassed by mutations in *flhA* and *flhB* that overcome the loss of the flagellar ATPase ^8,23,24^.

In addition to the FilJ-dependent gate opening signal, we have previously shown that a subunit N-terminal hydrophobic signal is required to trigger opening of the export gate^19^ (Fig. 1C). To determine whether both signals were required to trigger opening of the export gate or whether the signals were functionally redundant, we asked if suppressor mutations in the ABPQR export gate (FliR-F_113_A or FliR-G_117_D) that could compensate for loss of the subunit N-terminal hydrophobic signal could also overcome loss of the flagellar ATPase (and of the FilJ-dependent gate opening signal). We found that neither FliR-F_113_A nor FliR-G_117_D could suppress the export defect caused by the loss of FliJ-dependent activation of FlhA (Fig.4). Taking the reciprocal approach, we found that a mutation in *flhB* (P_28_T) that alleviates the loss of the FliJ-dependent activation signal could not overcome loss of the subunit N-terminal hydrophobic signal (Fig 4). This indicates that both signals are essential for efficient opening of the flagellar export gate. We propose that FliJ binding to FlhA and the presence of a subunit at the export machinery induce distinct conformational changes in FlhA, rendering the export machinery competent to utilise the pmf to energise opening of the export gate. The lack of either signal prevents opening of the export gate and, in turn, maintains the membrane permeability barrier (Fig. 5).

**Figure 6.**
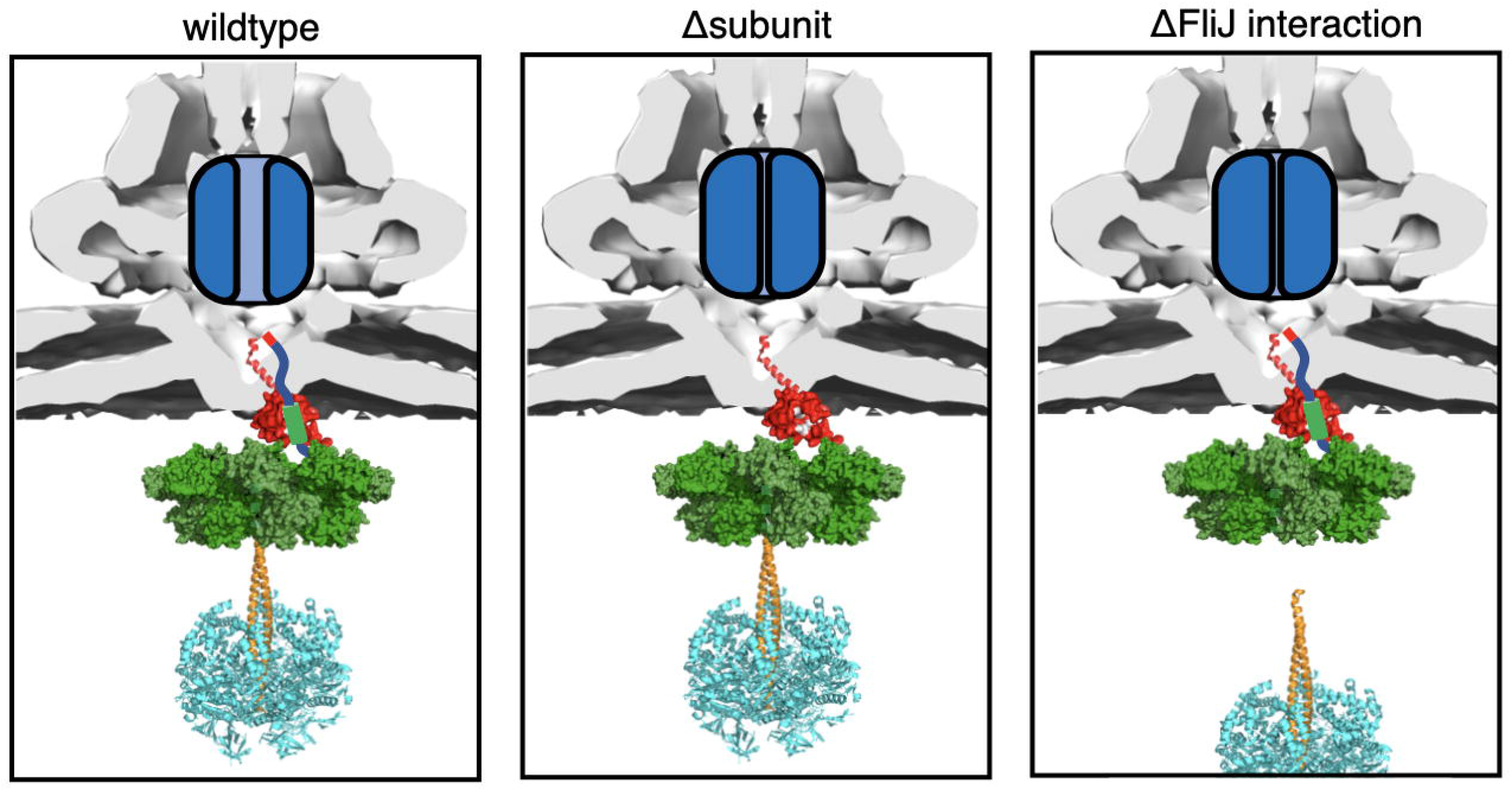
Two mutually exclusive signals are required to activate the export machinery and trigger opening of the export gate. A cartoon schematic illustrating the effect of either (i) the absence of a subunits docked at the export machinery (middle, Δsubunit) or (ii) the absence of a FliJ-FlhA interaction (right, ΔFliJ) (required to convert the export machinery into a highly efficient ΔΨ-driven export machine) on the ability of the export machinery to energise export gate opening (blue). ATP hydrolysis by the FliI (orange, PDB: 2DPY) component of the ATPase complex is thought to drive rotation of the FliJ stalk subunit (orange, PDB: 3AJW), allowing FliJ to bind all nine binding sites on the nonameric ring of FlhA (green, PDB: 3A5I), converting the export machinery into a highly efficient ΔΨ-driven export machine. Early flagellar subunits dock at FlhBc (red, PDB: 3B0Z) and contain a N-terminal hydrophobic export signal required for efficient subunit export and translocation through the export gate complex (left). The absence of subunits at the export machinery (middle) or the absence of FliJ-FlhA interactions (right) renders the export machinery unable to utilise the PMF to drive opening of the export gate.

Based on the structural and functional similarities between the core components of the injectosome and flagellar Type III Secretion Systems, it is highly probable that the mechanism of export gate opening is conserved. In summary, we propose that the pmf induces conformational changes in FlhA that are transmitted to the adjacent FliPQR-FlhB_N_ components of the export gate, energising gate opening. We demonstrate that FlhA requires at least two distinct signals to facilitate subunit export: a FliJ-dependent activation signal and a subunit docked at the export machinery. Our data suggest a multi-signal control system that acts *via* FlhA to ensure that use of the pmf and export gate opening only occurs when a FliJ activation signal is received and a subunit is available for export.

## Materials and Methods

### Bacterial strains, plasmids and growth conditions

*Salmonella* strains and plasmids used in this study are listed in table 1. The Δ*flgD*::K_m_^R^ strain in which the *flgD* gene was replaced by a kanamycin resistance cassette and the Δ*flgBCDEFGHIJ*::Km^R^ strain in which a large portion of the *flg* operon was replaced with a kanamycin cassette were constructed using the λ Red recombinase system^33,34^. Strains containing chromosomally encoded FliR and/or FlhA variants were constructed by aph-I-SceI Kanamycin resistance cassette replacement using pWRG730^34^. Recombinant proteins were expressed in *Salmonella* from isopropyl β-D-thiogalactoside-inducible (IPTG) inducible plasmid pTrc99a^35^. Bacteria were cultured at 30–37 °C in Luria-Bertani (LB) broth containing ampicillin (100 μg/ml).

### Choline sensitivity assay

The choline sensitivity assay was performed essentially as described in Ward *et al*.^21^. Cells were grown overnight at 32°C in terrific broth (TB) medium with shaking (200 PRM). The overnight cultures were diluted 1:50 in TB and grown at 32°C with shaking (200 RPM) to an A600 1.0. These cultures were diluted 1:50 in TB containing choline at concentrations as indicated in Figure 2. The cultures were grown for 4.5 hours at 32°C with shaking (200 RPM) and the A_600_ measured. The Miles and Misra method^36^ was used to determine the colony forming units for strains grown in 300 mM choline. 10-fold dilutions of culture were spotted (10 μl) onto LB agar plates and grown overnight at 37°C.

### Flagellar subunit export assay

*Salmonella* strains were cultured at 37 °C in LB broth containing ampicillin and IPTG to mid-log phase (OD_600nm_ 0.6-0.8). Cells were centrifuged (6000*g*, 3 min) and resuspended in fresh media and grown for a further 60 minutes at 37 °C. Cells were pelleted by centrifugation (16,000*g*, 5 min) and the supernatant passed through a 0.2 μm nitrocellulose filter. Proteins were precipitated with 10% trichloroacetic acid (TCA) and 1% Triton-X100 on ice for 1 hour, pelleted by centrifugation (16,000*g*, 10 min), washed with ice-cold acetone and resuspended in SDS-PAGE loading buffer (volumes calibrated according to cell densities). Fractions were analysed by immunoblotting. For the export assays of strains supplemented with arginine, cells were treated essentially as described above except cells were resuspended in fresh media supplemented with 20 mM arginine and grown for a further 30 minutes instead of 60 minutes.

### Motility assays

For swimming motility, cultures were grown in LB broth to A600nm 1. Two microliters of culture were inoculated into soft tryptone agar (0.3% agar, 10 g/L tryptone, 5g/L NaCl) containing ampicillin (100 μg/ml). Plates were incubated at 37 °C for between 4 and 6 hours unless otherwise stated. For swarming motility, one microliter of overnight culture grown in LB broth was inoculated onto tryptone agar plates (0.6% agar, 1% w/v tryptone, 0.5% w/v sodium chloride) containing appropriate antibiotics and inducing agents and supplemented with 0.3% glucose and incubated at 30 °C for 12-16 hours.

### Screen for motile suppressors from the flhA-K_203_A variant

Cells of the *Salmonella flhA*-K_203_A *strain* were cultured at 37 °C in LB broth to mid-log phase and inoculated into soft tryptone agar (0.3% agar, 10 g/L tryptone, 5g/L NaCl). Plates were incubated at 30 °C until motile ‘spurs’ appeared. Cells from the spurs were streaked to single colony and the *flhA* gene sequenced. New strains containing the *flhA*-K_203_A and suppressor mutations were constructed by aph-I-SceI λ Red recombination to confirm that the mutations within *flhA* were responsible for the motility suppressor phenotype^34^.

### Quantification and statistical analysis

Experiments were performed at least three times. Immunoblot were quantified using Image Studio Lite. The unpaired two-tailed Student’s *t*-test was used to determine *p*-values and significance was determined as \**p* < 0.05. Data are represented as mean ± standard error of the mean (SEM), unless otherwise specified and reported as biological replicates.

## Author contributions

O.J.B and G.M.F conceived and designed experiments. O.J.B conducted experiments. O.J.B analysed the data. O.J.B and G.M.F wrote the paper

## Acknowledgements

This work was funded by a grant from the Biotechnology and Biological Sciences Research Council (BB/M007197/1) to G.M.F, and a University of Cambridge John Lucas Walker studentship to O.J.B.

## Materials & Correspondence

Materials are available from the corresponding authors upon request.

## Supplementary Figures

**Figure S1**

Whole cell (cell) proteins from late exponential-phase cultures of recombinant *Salmonella flgD* null strains producing chromosomally-encoded FliR variants (F_110_A, F_113_A, F_110_A+F_113_A or G_117_D) or wild type FliR (FliR), and a pTrc99a plasmid-encoded FlgD subunit variant in which residues 9-32 were replaced with two repeats of the six amino acid sequence Gly-Ser-Thr-Asn-Ala-Ser (FlgD_short_). Proteins were separated by SDS (15%)-PAGE and analysed by immunoblotting with anti-FlgD polyclonal antisera. Apparent molecular weights are in kilodaltons (kDa).

**Figure S2**

**A**. Swimming (top panel; 0.25% soft tryptone agar) and swarming (bottom panels; 0.6% agar-tryptone with 0.5% glucose) motility of recombinant *Salmonella* strains producing chromosomally-encoded FliR variants (F_110_A, F_113_A, G_117_D or F_110_A+F_113_A) or SJW1103 wild type.

**B**. Whole cell (cell) and secreted (secreted) proteins from late exponential-phase cultures of recombinant *Salmonella* strains producing chromosomally-encoded FliR variants (F_110_A, F_113_A, G_117_D or F_110_A+F_113_A) or SJW1103 wild type. Proteins were separated by SDS (15%)-PAGE and analysed by immunoblotting with anti-FliC, anti-FliD, anti-FlgK, anti-FlgD, anti-FlhA and anti-FlgN polyclonal antisera. Apparent molecular weights are in kilodaltons (kDa).

**Figure S3**

Whole cell (cell) proteins from late exponential-phase cultures of recombinant *Salmonella* strains producing a chromosomally-encoded FliR variant in which residues 110-116 of the FliR plug is deleted (Δplug), a *Salmonella* strain deleted for the genes (*flgBCDEFGHIJ*) that encode most of the early flagellar rod and hook subunits (Δsubunits), a *Salmonella* strain that both produces the chromosomally-encoded FliRΔplug variant and is deleted for the genes (*flgBCDEFGHIJ*) that encode most of the early flagellar rod and hook subunits (ΔplugΔsubunits) and a SJW1103 wild type strain (FliR) were separated by SDS (15%)-PAGE and analysed by immunoblotting with anti-FlhA and anti-FlgD polyclonal antisera. Apparent molecular weights are in kilodaltons (kDa).

**Figure S4**

Whole cell (cell) and supernatant (secreted) proteins from late exponential-phase cultures of a strain deleted for two genes that encode arginine decarboxylase enzymes (Δ*speA* and Δ*adiA*, labelled FlhA) or a *Salmonella* Δ*speA-*Δ*adiA* strain producing a chromosomally-encoded FlhA-K_203_A variant (K_203_A). Cells were supplemented with (+) or not supplemented with (-) 20 mM arginine 30 minutes prior to collection of whole cell and culture supernatants. Proteins were separated by SDS (15%)-PAGE and analysed by immunoblotting with anti-FlgK, anti-FlgD, anti-FlhA or anti-FlgN polyclonal antisera. Apparent molecular weights are in kilodaltons (kDa).

**Figure S5**

Whole cell (cell) proteins from late exponential-phase cultures of recombinant *Salmonella flgD* null strains producing chromosomally-encoded wild type FlhB or its variants (FlhB-P_28_T). Both strains produced a pTrc99a plasmid-encoded FlgD subunit variant in which residues 9-32 were replaced with two repeats of the six amino acid sequence Gly-Ser-Thr-Asn-Ala-Ser (FlgD_short_). Proteins were separated by SDS (15%)-PAGE and analysed by immunoblotting with anti-FlgD polyclonal antisera. Apparent molecular weights are in kilodaltons (kDa).

