## Supplementary figures and images for "Regulation of bacterial Type III Secretion System export gate opening"

### Supplemental Figure 1

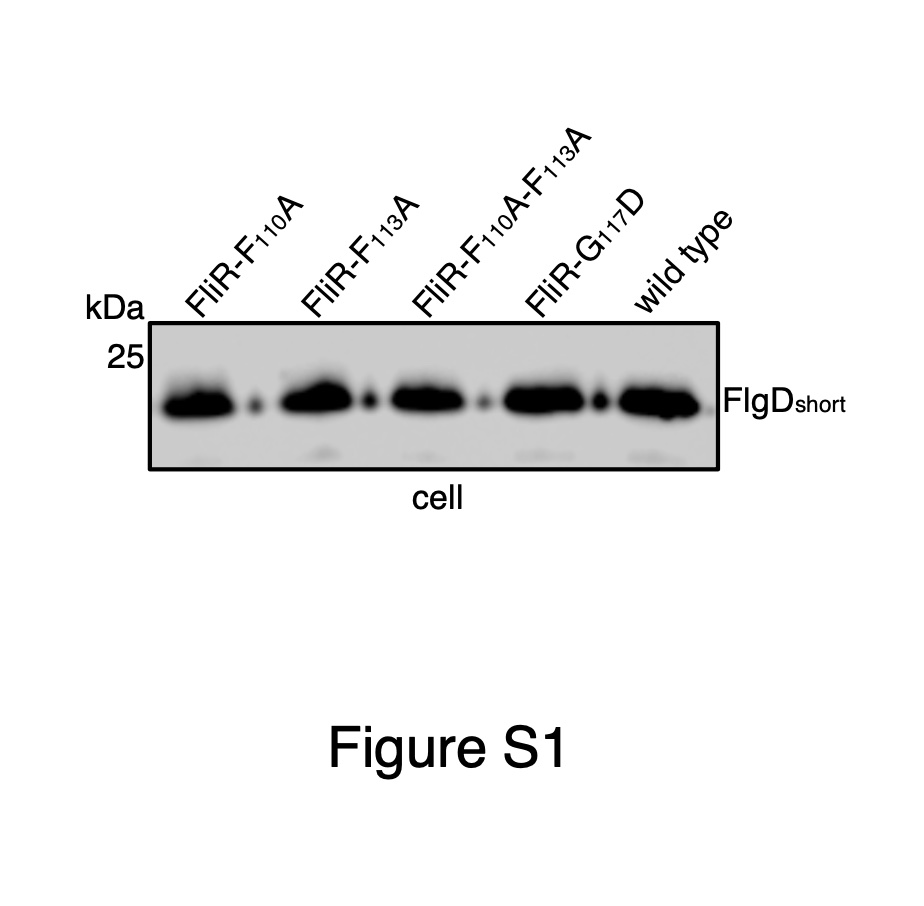

### Supplemental Figure 1

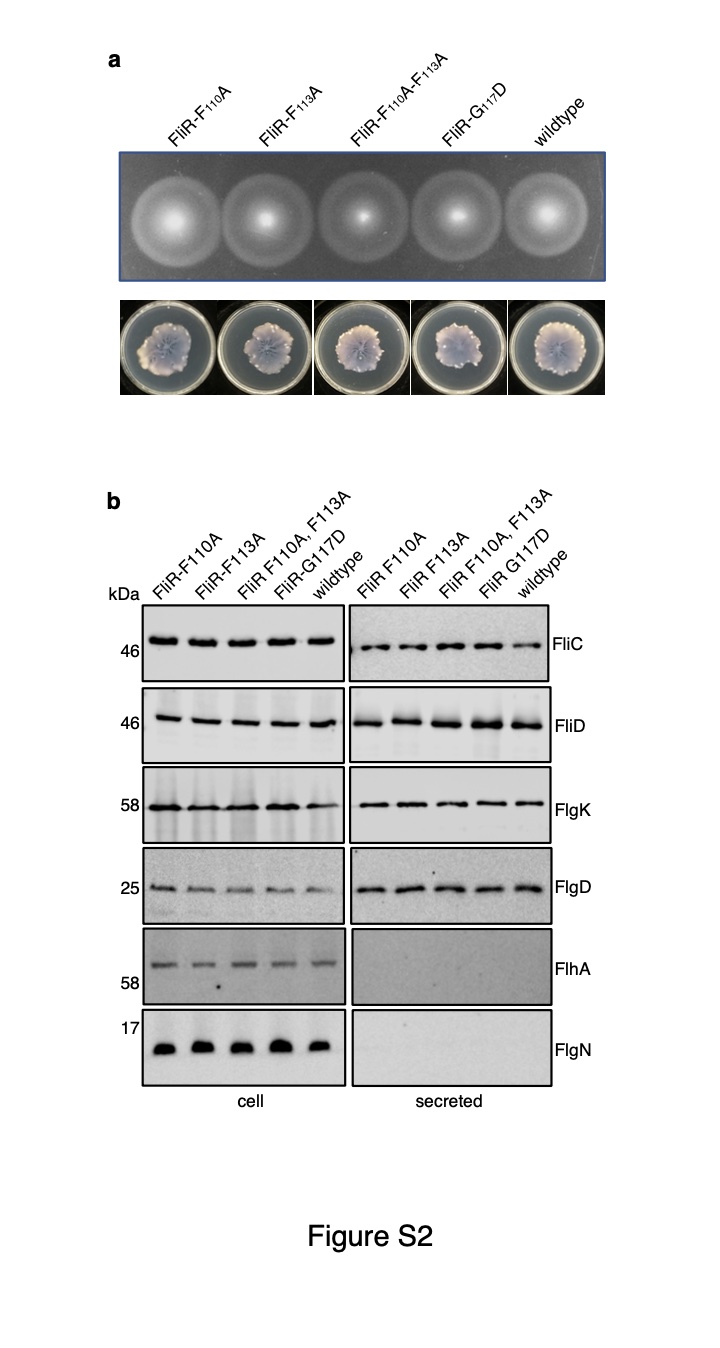

### Supplemental Figure 3

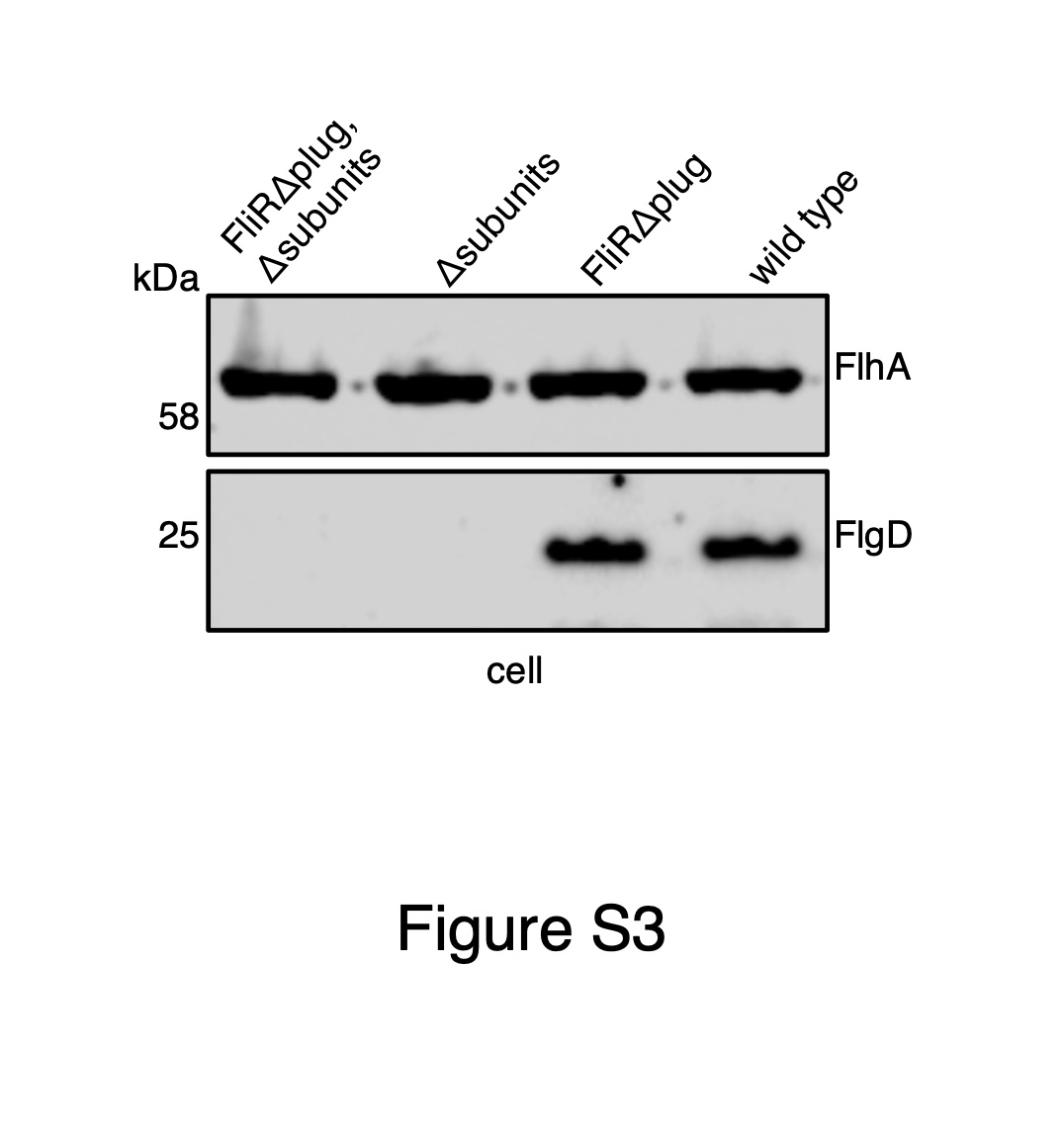

### Supplemental Figure 4

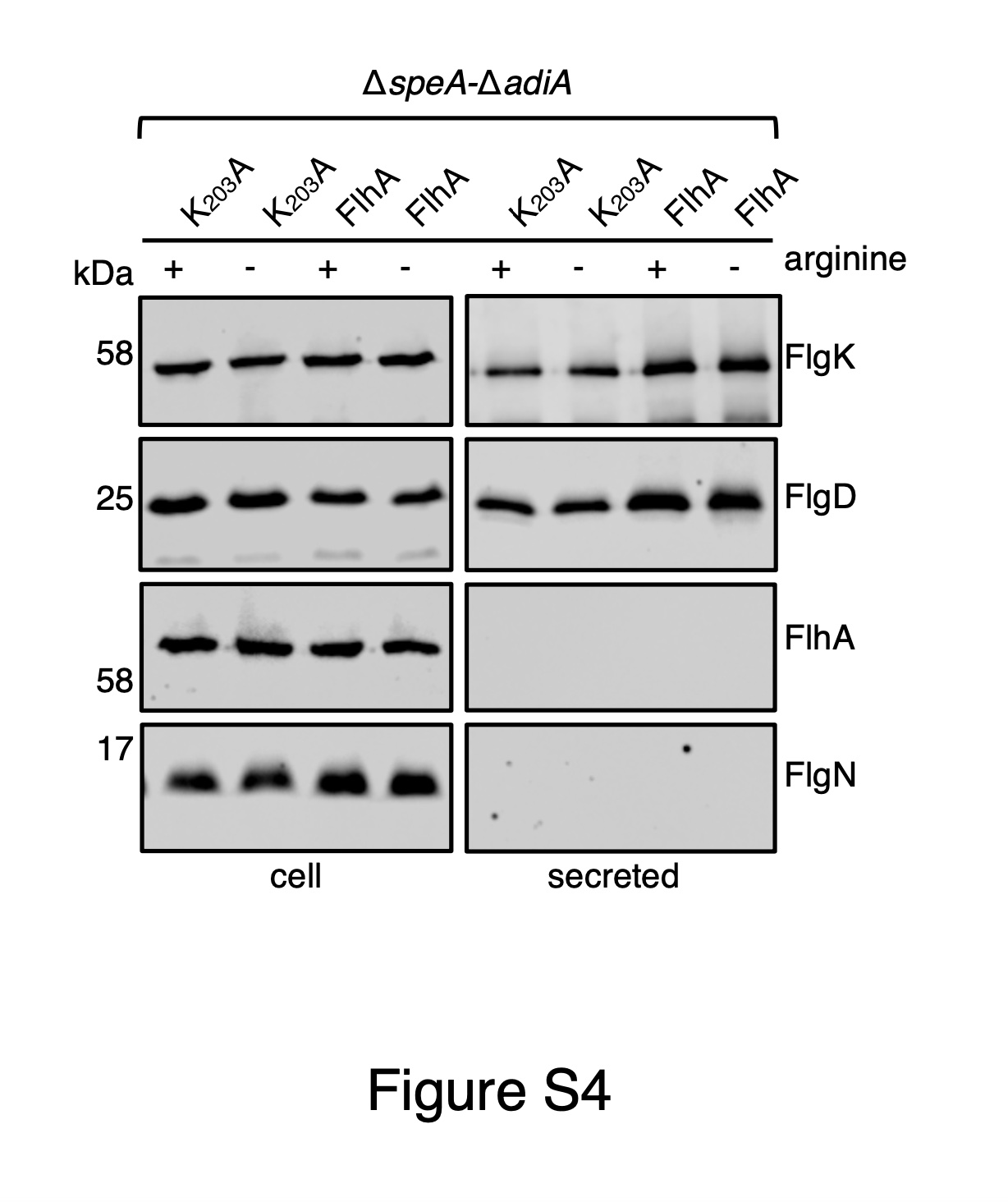

### Supplemental Figure 5

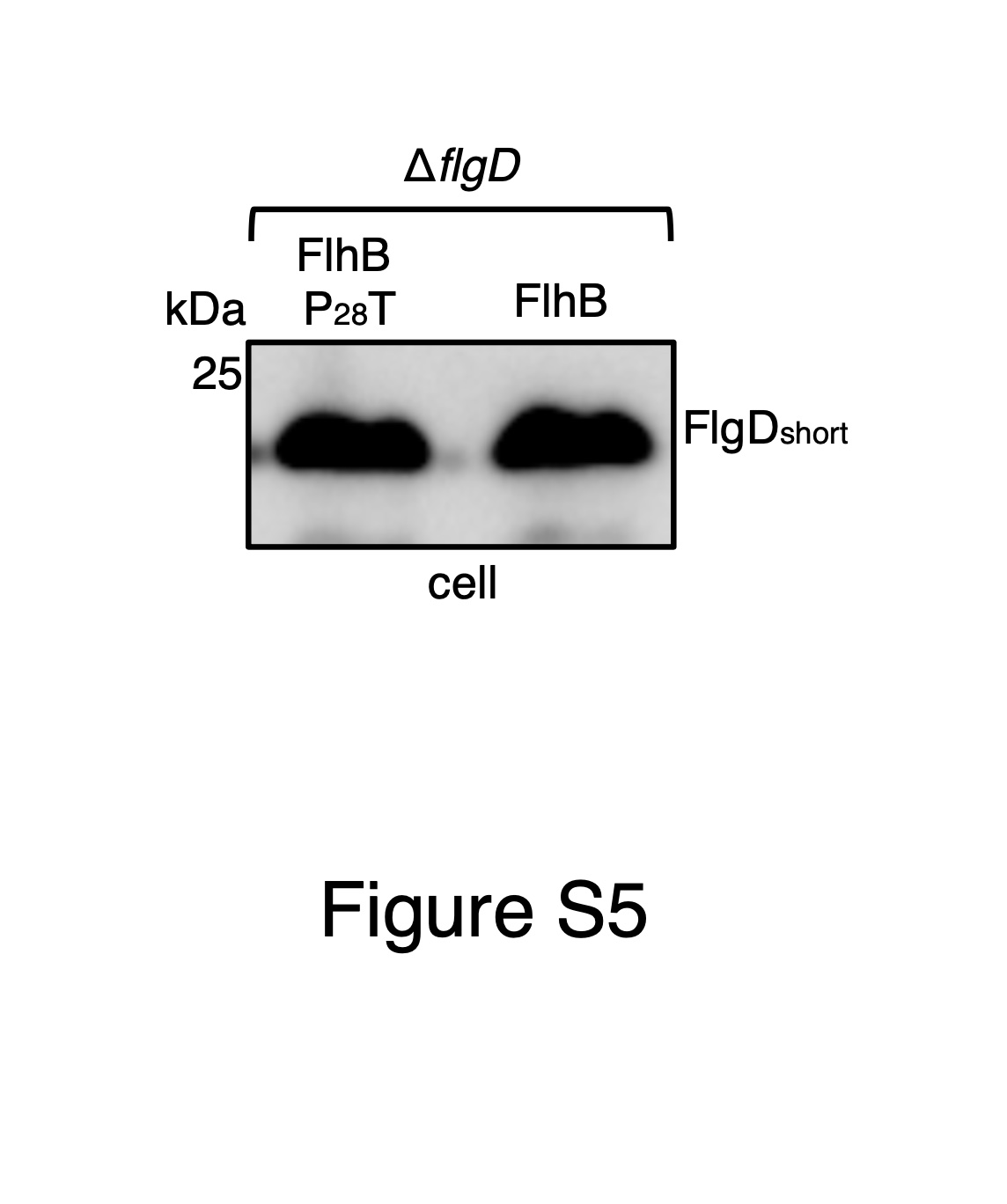
