## Supplemental Table 1 for "Regulation of bacterial Type III Secretion System export gate opening"

**Supplementary Table 1**. Strains and recombinant plasmids.

| Strains | Description |
|---|---|
| *Salmonella typhimurium* | |
| SJW1103 | wildtype |
| *flgD* null | Δ*flgD*::kmR |
| *fliR*(F110A) | *fliR* F110A variant |
| *fliR*(F113A) | *fliR* F113A variant |
| *fliR*(F110A,F113A) | *fliR* F110A,F113A variant |
| *fliR*(G117D) | *fliR* G117D variant |
| *fliR*(F110A), *flgD* null | *fliR* F110A variant, Δ*flgD*::kmR |
| *fliR*(F113A), *flgD* null | *fliR* F113A variant, Δ*flgD*::kmR |
| *fliR*(F110A,F113A), *flgD* null | *fliR* F110A, F113A variant, Δ*flgD*::kmR |
| *fliR*(G117D), *flgD* null | *fliR G117D* variant, Δ*flgD*::kmR |
| *fliR*(Δ110-116) | *fliR* Δ110-116 variant |
| *fliR*(Δ110-116), *flgBCDEFGHIJ null* | *fliR* Δ110-116 variant, Δ*flgBCDEFGHIJ*::KmR |
| *flgBCDEFGHIJ null* | Δ*flgBCDEFGHIJ*::K mR |
| *flhA*(K203A) | *flhA* K203A variant |
| *flhA*(K203A), *fliR*(F113A) | *flhA* K203A variant, *fliR* F113A variant |
| *flhA*(K203A), *fliR*(G117D) | *flhA* K203A variant, *fliR* G117D variant |
| *flhA*(K203A), *adiA* null, *speA* null | *flhA* K203A variant, Δ*adiA*::KmR, Δ*speA*::SpectR |
| *adiA* null, *speA* null | Δ*adiA*::KmR, Δ*speA*::SpectR |
| *fliR*(F113A), *fliHI* null, *flgM* null | *fliR* F113A variant, Δ*fliHI*, Δ*flgM*::SpectR |
| *fliR*(G117D), *fliHI* null, *flgM* null | *fliR* G117D variant, Δ*fliHI*, Δ*flgM*::SpectR |
| *flhB*(P28T), *fliHI* null, *flgM* null | *flhB P28T* variant, Δ*fliHI*, Δ*flgM*::SpectR |
| *fliHI* null, *flgM* null | Δ*fliHI*, Δ*flgM*::SpectR |
| *flhB*(P28T), *flgD* null | *flhB P28T* variant, Δ*flgD*::KmR |
| Plasmids | |
| pTrc99a FlgD | 1-232aa |
| pTrc99a $FlgD_{short}$ | 1-8, Gly, 2x(Gly- Ser-Thr-Asn-Ala- Ser), 33-232aa |
| pWRG730 | Lambda red recombinase under the P*L* promoter under the control of the temperature sensitive CI857 repressor. I-SceI under the Tet promoter. |
| pWRG717 | Kanamycin resistance cassette with I-*Sce*I recognition site |
